# Masked target visibility is selectively impaired by 20 Hz transcranial alternating current stimulation

**DOI:** 10.1101/2022.06.09.495501

**Authors:** Yayla Ilkson, Alethia de la Fuente, Enzo Tagliazucchi, Carla Pallavicini

## Abstract

It has been proposed that both conscious and unconscious perception are associated with a feedforward sweep of oscillatory activity in the gamma band (>40 Hz), while conscious perception also requires recurrent feedback via beta band (~20 Hz) oscillations. To investigate the causal relationship between these oscillations and (un)conscious visual perception, we assessed the effect of transcranial alternating current stimulation (tACS) in the gamma (40 Hz) and beta (20 Hz) bands on the objective and subjective visibility of targets in a metacontrast backward masking task. We expected that 40hz-tACS would affect both the correct categorization of the target (i.e. objective visibility) and the reports of conscious perception (i.e. subjective visibility). Moreover, we expected that 20Hz-tACS would selectively affect the subjective visibility. Our results showed that target visibility was selectively compromised by 20Hz-tACS but, in contrast to our hypothesis, this effect was specific to objective visibility. Although the power of local beta oscillations increased after 20Hz-tACS, inter-areal beta synchrony could have nevertheless been impaired, a possibility that should be investigated in the future by means of source reconstructed high density electroencephalography recordings. In summary, we provided evidence supporting that 20Hz-tACS is capable of modulating target visibility, suggesting a possible a causal link between synchrony in this frequency band and visual perception. Future studies could build upon this result by investigating other forms of stimulation and other model organisms, further contributing to our knowledge of how conscious access causally depends on brain oscillations.

## Introduction

Less than a century ago, the concept of consciousness was long considered a philosophic topic beyond the scope of empirical research. However, a surge of neuroscientific research on consciousness occurred in the last few decades, facilitated by new technologies and ingenious experimental paradigms designed to explore this elusive phenomenon (Crick and Koch 1990; Seth et al. 2008; Tagliazucchi 2020). A distinction is often made between research on conscious states (e.g. sleep, coma, anesthesia, and others) and research on conscious information access, which is related to the perceptual awareness of sensory inputs. Following the original proposal by Crick and Koch (Crick and Koch 1990), the majority of experiments designed to study the neural correlates of conscious perception are focused on vision, due to the rich nature of human visual experiences and the comparatively advanced knowledge of the primate visual system.

The influential neuronal global workspace (GW) theory equates consciousness with the global availability of information for its subsequence processing by multiple independent and parallel systems that implement specific cognitive functions (Dehaene et al. 2006). The GW theory provides a theoretical framework for the interpretation of neurophysiological observations related to conscious information access. According to this theory, the reportability of sensory percepts is the result of widespread communication (“broadcasting”) of information in the brain. This availability is manifest as the all-or-none and sustained ignition of frontal, parietal and temporal regions by stimuli that are both strong and attended (Dehaene and Naccache 2001; Baars 2002, 2005; Dehaene et al. 2006, 2011; Baars et al. 2013). Specifically, it is proposed that a reportable stimulus initially induces activations in the primary sensory areas, and then propagates to frontal, parietal and temporal higher-order associative areas, informing cognitive functions such as language production, decision making, motor planning and execution, among others. In contrast, unconscious (i.e. non reportable) stimuli induce weak activations that fail to propagate beyond early sensory regions, e.g. activations that remain limited to the visual cortex (Dehaene et al. 2006).The analysis of event-related potentials shows that unconscious visual processing occurs early (<270 ms) whereas conscious processing is related to later potentials (>270 ms), which are generated and sustained by a broad fronto-parieto-temporal network (Melloni et al. 2007; Del Cul et al. 2007). Alternatively, it is possible that strong but unattended stimuli become pre-conscious, in the sense that they are susceptible to ignite the global workspace and become globally available, but will not do so unless harnessed to the fronto-temporo-parietal network by redirecting top-bottom attention to them in a short window of time after their arrival to early sensory areas (Dehaene et al. 1998, 2001; Naccache and Dehaene 2001; van Gaal et al. 2014).

Lamme & Roelfsema (2000) proposed a slightly different view, advocating that recurrent feedback, not only feedforward processing, is crucial for the emergence of conscious awareness. They hypothesized that conscious and unconscious processing are associated with attended global and unattended local fast feedforward activity, respectively. Also, they proposed that unattended feedforward activity can extend beyond sensory areas and modulate behavior without awareness, consistent with the GW theory notion of pre-conscious states (Dehaene et al. 1998, 2001; Naccache and Dehaene 2001; van Gaal et al. 2014). This can be illustrated by the well-known phenomenon of blindsight, i.e. the statistically correct categorization of visual stimuli in the absence of reports of conscious perception. The main difference between conscious and unconscious processing, according to the hypothesis by Lamme and Roelfsema, is that conscious processing requires recurrent feedback processing, which might be important for the top-bottom amplification of the incoming sensory signals (Sergent and Dehaene, 2004).

Gamma oscillations have often been associated with conscious information access (Sergent and Dehaene 2004; Melloni et al. 2007; Steinmann et al. 2014; Cavinato et al. 2015). However, instead of gamma synchrony, Gaillard et al. (2009) observed increased beta synchrony during unmasked vs, masked stimuli presentation, measured in epileptic patients with intracranial EEG. Moreover, this activity pattern was restricted to late processing (300-500 ms) and was recurrent in the case of conscious information access, in line with the proposal by Roelfsema & Lamme. Similarly, Bastos et al. (2015) showed that feedforward activity in the visual cortex is carried by gamma oscillations, while feedback influences depend on beta oscillations. Interestingly, microstimulation of lower visual areas in monkeys induces gamma-oscillations in higher visual areas, while communication in the opposite direction was governed by alpha frequencies (van Kerkoerle et al. 2014).

Overall, the aforementioned studies suggest that low-level subliminal or pre-conscious information processing can occur during a fast feedforward sweep, involving local activity in the gamma range that quickly fades away. In contrast, conscious processing requires feedback influences of distant brain areas carried by beta oscillations. Importantly, most of the current findings supporting this scenario are either correlational or have been obtained in animal models; for instance, it is not known whether disrupting feedback in the beta band selectively impairs conscious access. Non-invasive electrical stimulation is a suitable method to investigate this causal link in humans. In particular, transcranial alternating current stimulation (tACS) applies a sinusoidal current in a chosen frequency and is capable of modulating neural oscillations at the stimulation frequency. We applied tACS with stimulation frequencies in the gamma (40 Hz) and beta (20 Hz) bands to participants performing a metacontrast backward-masking task. We expected that gamma-tACS would affect the correct categorization of unreported targets (i.e. objective visibility) and the explicit report of the conscious perception of the targets (i.e. subjective visibility); in contrast, we hypothesized that beta-tACS would impact only on the subjective visibility, as it would interfere with the recurrent synchronization implicated with conscious access.

## Methods

### Participants

This study included 34 right handed healthy participants. Three subjects did not complete all the experimental sessions and were excluded, resulting in a total of 31 subjects (8 women and 23 men; mean age: 30.39 ± 5.13, range age: 22-41). All participants and researchers were blind with respect to the stimulation condition applied in each session. All participants had normal or corrected-to-normal eye vision, no history of neurological disease and did not receive anti-epileptic treatment. Furthermore, subjects did not undergo treatment with psychotropic drugs that affected cortical excitation, nor had a pacemaker, cardiac defibrillator implants, nor intracranial brain stimulation electrode implants. All participants gave written informed consent in accordance with the Helsinki declaration. This study was approved by the ethics committee of Hospital General de Agudos José María Ramos Mejía in the city of Buenos Aires, Argentina, and conducted in accordance with international principles for human medical research, including the declaration of Helsinki. Subjects did not receive compensation for their participation in this study.

### Experimental task

A metacontrast backward-masking (Del Cul et al. 2007) paradigm was implemented in PsychoPy v3.0 (Peirce et al. 2019) and performed by the participants during the different stimulation conditions. In each trial of this task, a target stimulus (consisting of the numerals 2, 3, 7 or 8) was followed by a mask with a variable temporal separation from the target, known as stimulus onset asynchrony (SOA; see Figure 1A). Both mask and targets were randomly presented either to the left or to the right of a fixation cross on the 13 inch display of a MacBook Pro Retina (late 2013 model) with a refresh rate of 60 Hz. Following the presentation of the masked target, two forced-choice questions were presented in the screen. First, participants had to indicate whether the target was larger or smaller in magnitude than the numeral 5, thus providing an objective visibility measure. Second, participants had to report whether they saw the target numeral, which constitutes a subjective visibility measure. The complete task consisted of 400 trials with evenly distributed SOAs (16.7, 33.3, 50, 66.7 and 83.3 ms). Sixty randomly distributed trials lacked targets before the mask to control for subject engagement.

**Figure 1:**
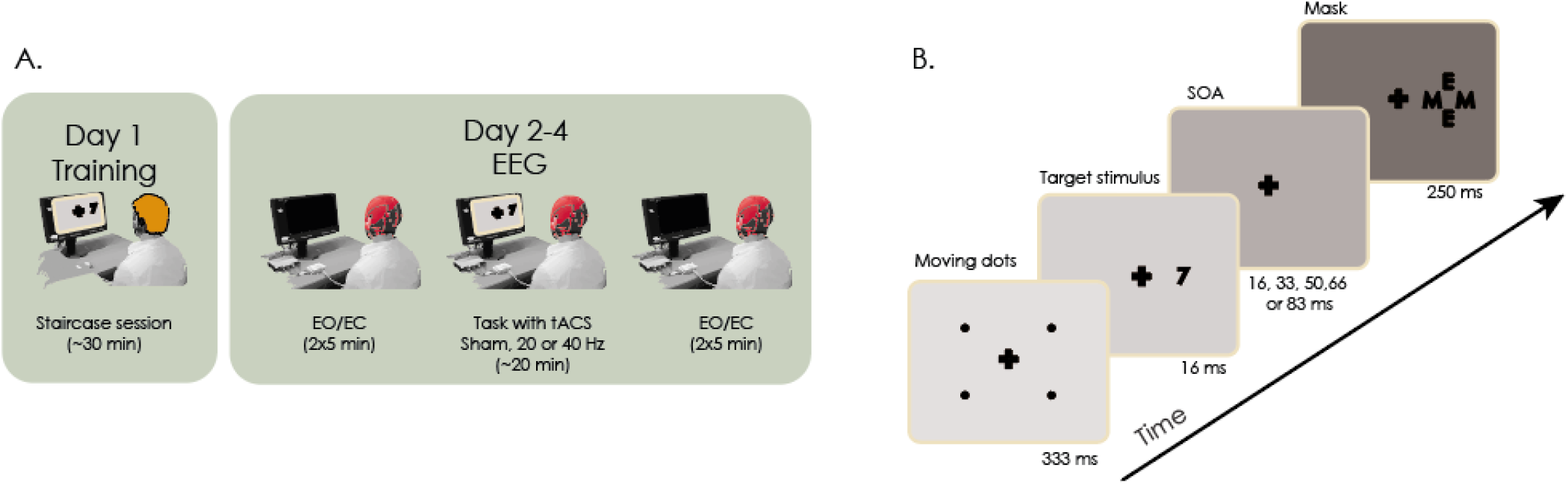
Task set-up and experimental procedure. **a)** In total, participants performed four sessions on four non-consecutive days. The first session included a staircase procedure to determine the appropriate target contrast for that participant. The remaining three sessions were identical in terms of the performed task, but were conducted under different tACS stimulation conditions (sham, 20Hz or 40Hz). Resting state EEG (eyes open and closed) was recorded for five minutes before and after stimulation. At the end of each stimulation session, participants completed a questionnaire to evaluate the somatosensory effects induced by tACS. **b)** Each trial of the backward masking task began with a set of four points moving to draw the attention of the participant to the fixation cross. Next, the target numeral briefly appeared in the display and was followed by a mask appearing at the past location of the target. The separations between target and mask (SOA) were uniformly spaced and randomly changed across trials.

### Determination of the subjective visibility threshold

A staircase procedure was implemented to determine a stimulus-background contrast for which each participant achieved subjective visibility in ~50% of the trials at the SOA of 50 ms. The contrast of the numeral presented as the masked stimulus could linearly range from −1 (gray, background color) to 1 (black). At the beginning of the staircase session, the contrast was set to 0.7 and could increase or decrease in steps of 0.2, depending of the response given by the participant at each trial. The contrast decreased when the objective response was correct and the subjective visibility rating was affirmative, and increased when the subject either incorrectly compared the stimulus to 5, or declared absence of subjective visibility. The final contrast used for the experimental task was computed as the mean of the last eight reversals of the staircase session.

### Protocol for tACS stimulation

Beta- (20Hz), gamma- (40hz) and sham-tACS were applied using two hybrid tCS and EEG electrodes with a circular Ag/AgCl contact area (diameter: 12 mm; 8-channel Neuroelectrics STARSTIM dual EEG-tACS). The electrodes were placed at the Oz and Fz locations of the standard10-10 EEG system. An intensity of 1000 μA (2000 μA peak-to-peak) with a ramp-up of 20 seconds was applied. During sham-tACS, a frequency of 30Hz was applied and discontinued after the ramp-up phase. Participants underwent tACS while they performed the metacontrast backward-masking task (duration ~20 min). The stimulation conditions were randomized and uniformly distributed throughout sessions and across participants. The randomization was performed by an individual not involved with the experiment, and the order of the stimulation conditions was stored in a spreadsheet that could be accessed by the researchers only at the data analysis step; thus, both participants and researchers were blind with respect to the applied stimulation.

### EEG recording

Complete EEG recordings were obtained for half of the participants (n=17). Resting state EEG was recorded preceding and following each tACS session with an 8-channel Neuroelectrics STARSTIM dual EEG-tACS. In addition to the electrodes located at the stimulation sites (Fz and Oz), EEG signals were recorded at T3 and T4 (left/right temporal areas), P3 and P4 (left/right centro-parietal areas) and F7 and F8 (left/right frontal areas) using Ag/AgCl electrodes placed according to the 10-10 EEG system. Ground and reference electrodes were placed on the earlobe. Data was sampled at 500 Hz with 24 bits AD conversion and streamed via Bluetooth to a notebook.

### Experimental sessions

Participants performed a total of four sessions on four non-consecutive days (average days between sessions: 6.14 ± 3.15, between 1 and 15 days). On the first session, participants performed a short training followed by the staircase. On the second, third and fourth sessions participants performed the metacontrast backward-masking task, while being stimulated with either 20Hz-, 40Hz or sham-tACS (counterbalanced and double-blind). Furthermore, preceding and following each tACS session, resting state EEG was recorded for 5 min with eyes closed (EC-EEG) and 5 min with eyes open (EO-EEG; see Figure 1B) for a subset of the participants. Participants performed the sessions approximately at the same time of the day as in their previous sessions (average hour difference between sessions: 2:03). All experimental sessions were performed in a darkened and acoustically isolated room. At the end of each session with tACS stimulation, subjects completed a standardized questionnaire (Antal et al. 2017) to assess the somatosensory effects elicited by the stimulation (e.g. itching, burning, etc).

### Statistical analysis of task performance

Statistical analyses were conducted using the R programming language (R Core Team 2014). Objective and subjective visibility were assessed based on the comparison with numeral 5 (‘correct’ / ‘incorrect’), and on the answer received when asked about the perception of the target number (‘seen’ / ‘unseen’). The blindsight rate was computed as the proportion of trials with the combination of ‘correct’ and ‘unseen’. The sightblind rate (name chosen to reflect the opposite of blindsight) was computed as the proportion of trials with the combination of ‘incorrect’ and ‘seen’. Each response measure consisted of a vector of binary outputs (‘correct’ / ‘incorrect’, ‘seen’/’unseen’). Mixed logistic regressions were used to test for conditional differences in the objective and subjective visibility rates (Bliksted et al. 2017; Chang et al. 2017; Łukowska et al. 2018). Each response measure was modeled as a function of tACS condition (categorical), session number (categorical) and SOA (continuous); subject IDs, target locations (left or right) and target numerals were included as random effects. Residuals were visualized as a function of the fitted values to check for nonlinearity.

Likelihood ratio tests between the full models (as described above) and the equivalent model reduced by tACS condition were used to statistically test for the effects of tACS. Additionally, to evaluate whether an interaction between the tACS condition and the SOA should be included into the model, an interaction model was compared to a model without interactions. Subsequently, to test the statistical significance of the coefficients of each model, confidence intervals were obtained and p-values were provided by either an ANOVA (for the logistic regressions) or t-tests (for the linear models) via a Satterthwaite’s degrees of freedom method. An alpha level of 0.05 was used for all tests. To provide further characterization of the task performance, the area under the curve (AUC) of the objective visibility vs. SOA was computed. The AUC was approximated via the trapezoidal method for the objective and subjective visibility as a function of SOA. This approximation was calculated by the following formula:

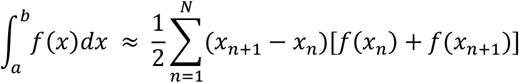

where *a* = *x*_1_, < *x*_2_ < … < *x_N_* < *x*_*N*+1_ = *b* and *x*_*n*+1_ – *x_n_* is the spacing between each consecutive pair of points. Conditional differences in AUC were tested with a linear model; specifically, tACS condition and session number were included as fixed effects and subject IDs as random effects.

### Questionnaires of somatosensory effects

The questionnaire ratings of potential somatosensory effects induced by tACS were normalized to scores ranging from zero to three for each sensation, with zero corresponding to feeling nothing, and three to feeling the sensation with strong intensity. These numerals were averaged for each subject to obtain one score for each condition. A linear model was used to describe the relationship between these scores and the tACS condition; specifically, tACS and session numbers were included as fixed effects and subject IDs as random effects.

### EEG data analysis

EEG pre-processing steps were performed using the EEGLAB Matlab toolbox (2018). EEG signals were band-pass filtered between 1-90 Hz and notch filtered between 47.5-52.5 Hz. Flatline channels were rejected (based on standard deviations 10 times lower than the other channels). The ‘clean_rawdata’ EEGLAB plugin (https://github.com/sccn/clean_rawdata) was used with default settings to correct continuous EEG data and reject bad channels using artifact subspace reconstruction. Subsequently, channels were interpolated, re-referenced to an average reference, and cut in epochs of two seconds. First, an amplitude threshold of −500 and 500 μV was applied to reject epochs with major artifacts. Next, noisy epochs were detected and rejected using a six standard deviation limit for single channels and a two standard deviation limit for all channels. Independent component analysis was performed to remove eyeblinks and other remaining sources of noise. To control for rank deficiency, the number of independent components was set to the number of channels that remained after channel rejection (i.e. eight channels – rejected channels) using principal component analysis.

### EEG spectral analysis

The power spectra of baseline EEGs (i.e. recorded before tACS application) were subtracted from those of the post tACS EEG measurements. Spectral power differences at 20Hz and 40Hz were computed separately for the eyes open and eyes closed EEG datasets, resulting in four comparisons (i.e. 20Hz with eyes open, 20Hz with eyes closed, 40Hz with eyes open, 40Hz with eyes closed). For each comparison, a one-way repeated measures ANOVA with tACS condition as a factor was performed for all eight channels. Benjamini-Hochberg’s False Discovery Rate (FDR) was employed to correct for multiple comparisons, with alpha level set to 0.05. Whenever significant results were found, post-hoc paired t-tests were performed to determine which conditions showed significant spectral power differences.

## Results

### Subjective and objective visibility as a function of SOA

Figure 2A and 2B show the objective and subjective visibility ratings as a function of SOA for all stimulation conditions. It is clear that both visibility measures increase with SOA. Indeed, SOA significantly affected the objective (*Z* = 44.18, *p* = 2e-16) and subjective (*Z* = 75.08, *p* = 2e-16) visibility. Specifically, for each unitary increase in SOA the odds of a ‘correct’ and a ‘seen’ trial changed 1.78 (95% CI: 1.73, 1.82) and 2.69 (95% CI: 2.62, 2.76) for objective visibility and subjective visibility, respectively. Even though we performed a staircase session to determine an appropriate personalized contrast at which ~50% of the targets were visible for a SOA of 50 ms, Figure 2B shows that the 50% subjective visibility threshold (averaged across participants) lies in the proximity of a SOA of 33.3 ms. Furthermore, the shape of the target visibility as a function of SOA did not exactly match previously reported findings (see supplementary figure 1 of Del Cul et al. 2007) since instead of a sudden increase in visibility, our data presented a more gradual increase (Figure 2A).

**Figure 2:**
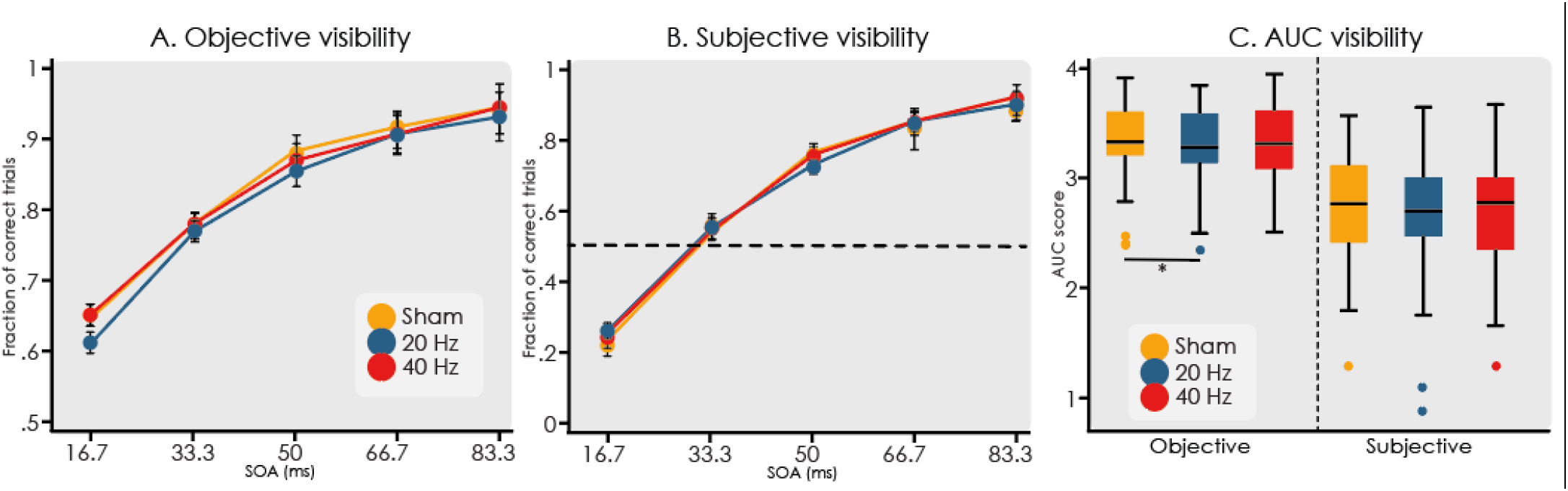
Objective and subjective. visibility between tACS conditions. The target visibility rate as a function of SOA with error bars representing standard errors of the rates. **a)** The objective visibility curve during 20Hz-tACS was significantly lower compared to sham-tACS (Z = −4.16, p = 3.17e-05). No difference between 40Hz-tACS and sham-tACS was found (Z =-1.61, p = 0.12). **b)** The striped line shows that the subjective visibility threshold lies slightly below a SOA of 33.3 ms. No differences between tACS conditions were found for subjective visibility (χ2(2) = 0.7, p = 0.7). **c)** The AUC scores for each tACS condition for the objective and subjective visibility curves. For the objective visibility, the AUC value for 20Hz-tACS is significantly lower than the value obtained for sham-tACS (t = 2.42, p = 0.018).

### Effects of tACS stimulation

In order to fully characterize the response, we performed likelihood ratio tests to assess the variables included in the mixed logistic model (see methods). For the objective visibility rates, this test showed that the interaction between tACS condition and SOA was not significant (χ2(2) = 0.3, *p* = 0.86) and therefore was not included in the final model. An additional likelihood ratio test showed that tACS condition significantly affected the objective visibility rates (χ2(2) = 17.63, *p* = 0.00015) and the odds ratio of a ‘correct’ trial for 20Hz-tACS compared to sham-tACS was 0.84 (95% CI: 0.78, 0.91) (ANOVA with Satterthwaite’s method, *Z* = −4.16, *p* = 3.17e-05). In other words, the probability of a ‘correct’ trial decreased during 20Hz-tACS compared to sham-tACS. Furthermore, the odds ratio of a ‘correct’ trial for 40Hz-tACS compared to sham-tACS was not significant (odds ratio = 0.94, 95% CI: 0.86, 1.01;*Z* =-1.61, *p* = 0.12). The AUC of the objective visibility as response to SOA was computed for each tACS condition (Figure 2C). A one-sided likelihood ratio test showed that tACS condition significantly affected the AUC (χ2(2) = 5.67, *p* = 0.029). In line with the previous results, AUC decreased during 20Hz-tACS compared to sham-tACS (decreased by −0.073, 95% CI: −0.13, −0.013, *t* = −2.42, *p* = 0.018).

For the subjective visibility rates (Figure 2B), the data showed no significant interaction between tACS condition and SOA (χ2(2) = 5.57, *p* = 0.062) and therefore this interaction was not included in the final model. Moreover, an additional likelihood ratio test showed that tACS condition did not significantly affect the subjective visibility rates (χ2(2) = 0.7, *p* = 0.7). Consistently, tACS condition did not affect the computed AUC for the subjective visibility as response to SOA (χ2(2) = 0.23, *p* = 0.89), as shown in Figure 2C.

To determine whether the decrease in objective visibility reflected a decrease in subliminal processing, we additionally analyzed blindsight rates (Figure 3A). In this case, the interaction between tACS condition and SOA was not significant (χ2(2) = 0.91, *p* = 0.63) and therefore not included in the final model. The final model showed that the odds of a blindsight trial significantly increased with 1.14 (95% CI: 1.1, 1.18) for each unitary increase in SOA (*Z* = 7.211, *p* = 5.53e-13; see figure 3A). Moreover, the data showed that tACS condition significantly affected the blindsight rate (χ2(2) = 6.79, *p* = 0.034). Specifically, the odds of a blindsight trial for 20Hz-tACS compared to sham-tACS was 0.88 (95% CI: 0.79, 0.97), or in other words, the probability of a blindsight trial decreased during 20Hz-tACS (*Z* = 2.57, *p* = 0.01). The 40Hz-tACS coefficient was not significant, meaning that there was no difference in blindsight rates between 40Hz-tACS and sham-tACS (odd ratio: 0.96, 95% CI: 0.86, 1.06), *Z* = −0.89, *p* = 0.38).

**Figure 3:**
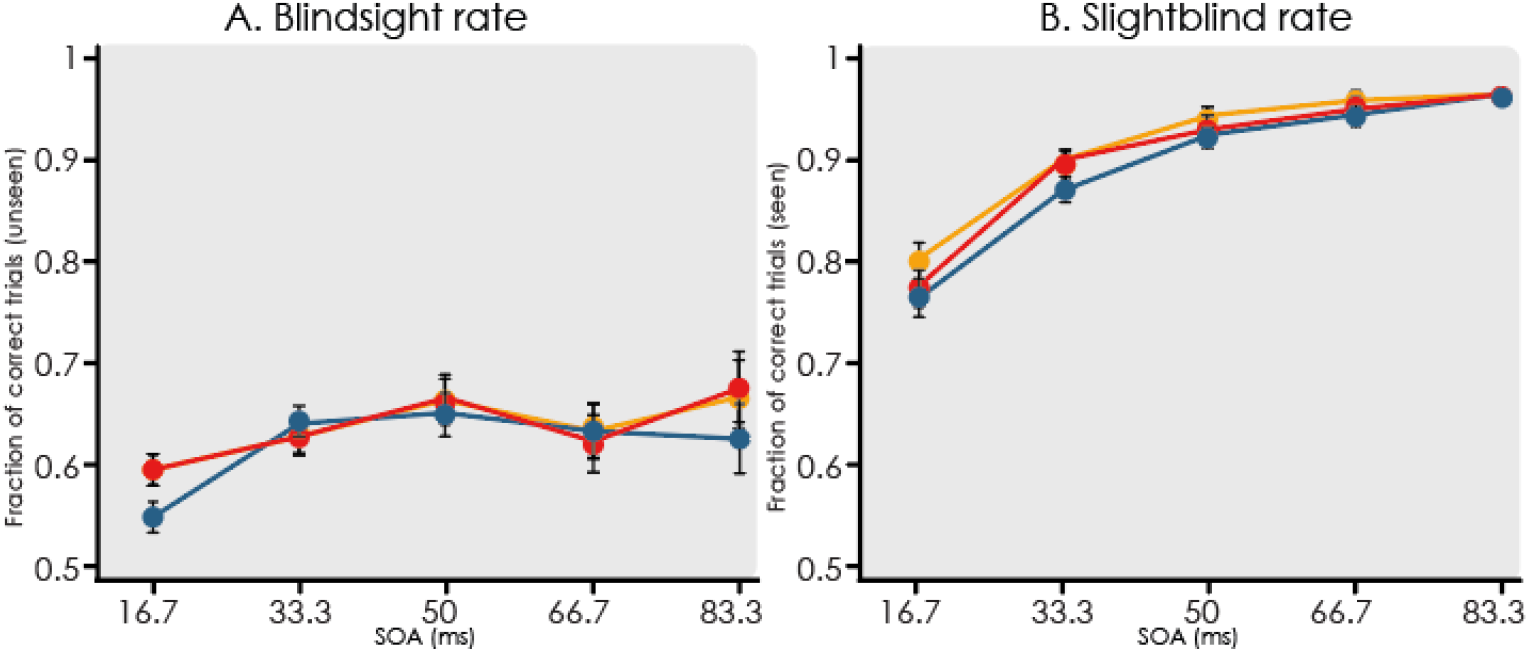
**a)** The blindsight rate for 20Hz-tACS was lower compared to sham-tACS (Z = 2.57, p = 0.01). **b)** The sightblind rates of both 20Hz-tACS (Z = −3.6, p < 0.001) and 40Hz-tACS (*Z* = −2.22, *p* < 0.05). were lower compared to sham-tACS.

The sightblind rate was analyzed to verify whether the decrease in blindsight rate actually represented a decrease in subliminal processing or was a result of a general decrease in objective visibility (Figure 3B). The data showed that tACS condition also affected sightblind (χ2(2) = 13.11, *p* = 0.0014), with an odds ratio of 0.75 (95% CI: 0.64, 0.86) for ‘correct; and ‘seen’ trials during 20Hz-tACS compared to sham-tACS (*Z* = −3.6, *p* = 0.00032). This result suggests that the decrease in blindsight is not specific to subliminal processing but an effect of a general decrease in objective visibility. Interestingly, the odds ratio of a sightblind trial during 40Hz-tACS was 0.83 (95% CI: 0.71, 0.98) and significant as well (*Z* = −2.22, *p* = 0.026). Moreover, similarly to the previous results, SOA significantly affected sightblind (odd ratio: 0.78, 95% CI: 1.69, 1.87, *Z* = 22.97, *p* = 2e16)

The reaction times (RT) for the objective response were analyzed to determine whether they were related to the decrease in objective visibility. Pearson’s correlations between the proportion of correct trials (objective visibility) and RTs were performed; however, no significant correlations were found (*t*(91) = 1.09, *p* = 0.28).

Next, we analyzed the questionnaire ratings to determine whether subjects were able to distinguish between tACS conditions based on the somatosensory perceptions they experienced. Interestingly, tACS condition significantly affected the somatosensory scores (χ2(2) = 9.22, *p* = 0.01), increasing the total score by 0.11 (95% CI: 0.03, 0.33) during 20Hz-tACS compared to sham-tACS (*t* = 2.721, *p* = 0.0085, Figure 4A). In contrast, no difference in scores between sham-tACS and 40Hz-tACS was found (*t* = 0.009, *p* = 0.99). Since 20Hz-tACS decreased the objective visibility as well as increased the questionnaire scores, a Pearson’s correlation test between these response measures was performed to investigate a possible relationship between both effects; however, no correlation was found (*t*(91) = 1.09, *p* = 0.31, Figure 4B).

**Figure 4:**
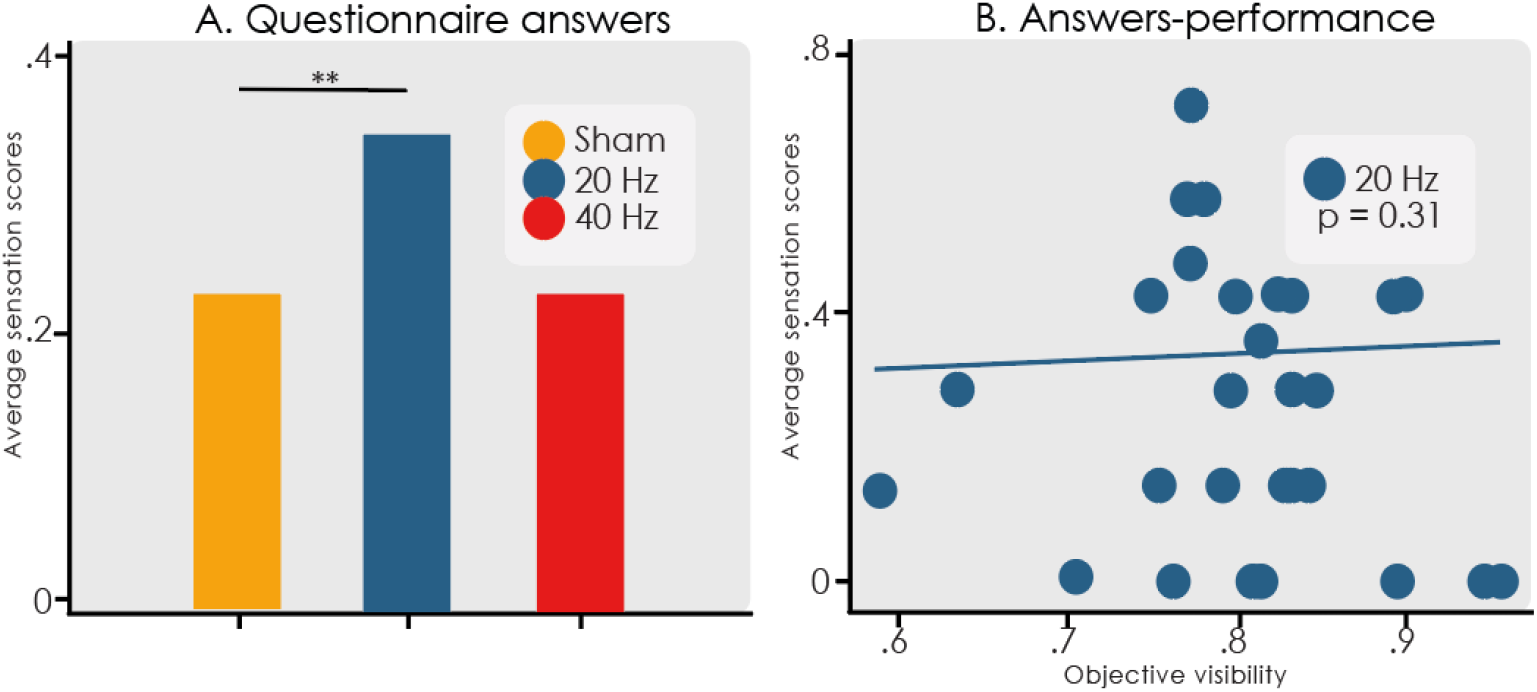
Questionnaire scores. a) Average somatosensory sensation scores per stimulation condition (**p<0.01). b) Scatter plot of average somatosensory sensation scores vs. objective visibility. No significant correlation was found between these two variables.

### EEG spectral power

Figures 5 and 6 present the differences in 20 Hz and 40 Hz power for each tACS condition, respectively, before and after the metacontrast backward-masking task. The topographic plots suggest that 20Hz power increased during the eyes closed condition after 20Hz-tACS (see Figure 5A). Indeed, channel F7 showed a significant power difference at 20Hz (*F*(2,15) = 6.43, *p* = 0.0029, FDR corrected) (Figure 5C). Both the 20Hz-tACS (*t*(16) = −3.17, *p* = 0.048) and 40Hz-tACS (*t*(16) = −3.17, *p* = 0.029) conditions increased 20 Hz power compared to sham-tACS (see Figure 5B). No other channels increased 20Hz power during the eyes closed condition, although Oz showed a tendency towards significance (*F*(2,15) =4.99, *p* = 0.0524) (Figure 5D). The 20 Hz power differences during the eyes open condition were less pronounced (Figure 5A). Only after the 40Hz-tACS condition a decrease in 20Hz spectral power was apparent; however, no statistically significant differences were found for 20 Hz power during eyes open. Figure 6 presents the differences in 40Hz power between tACS conditions. No significant differences were found in 40Hz power after 40Hz-tACS with both the eyes open and closed (Figure 6B)

**Figure 5:**
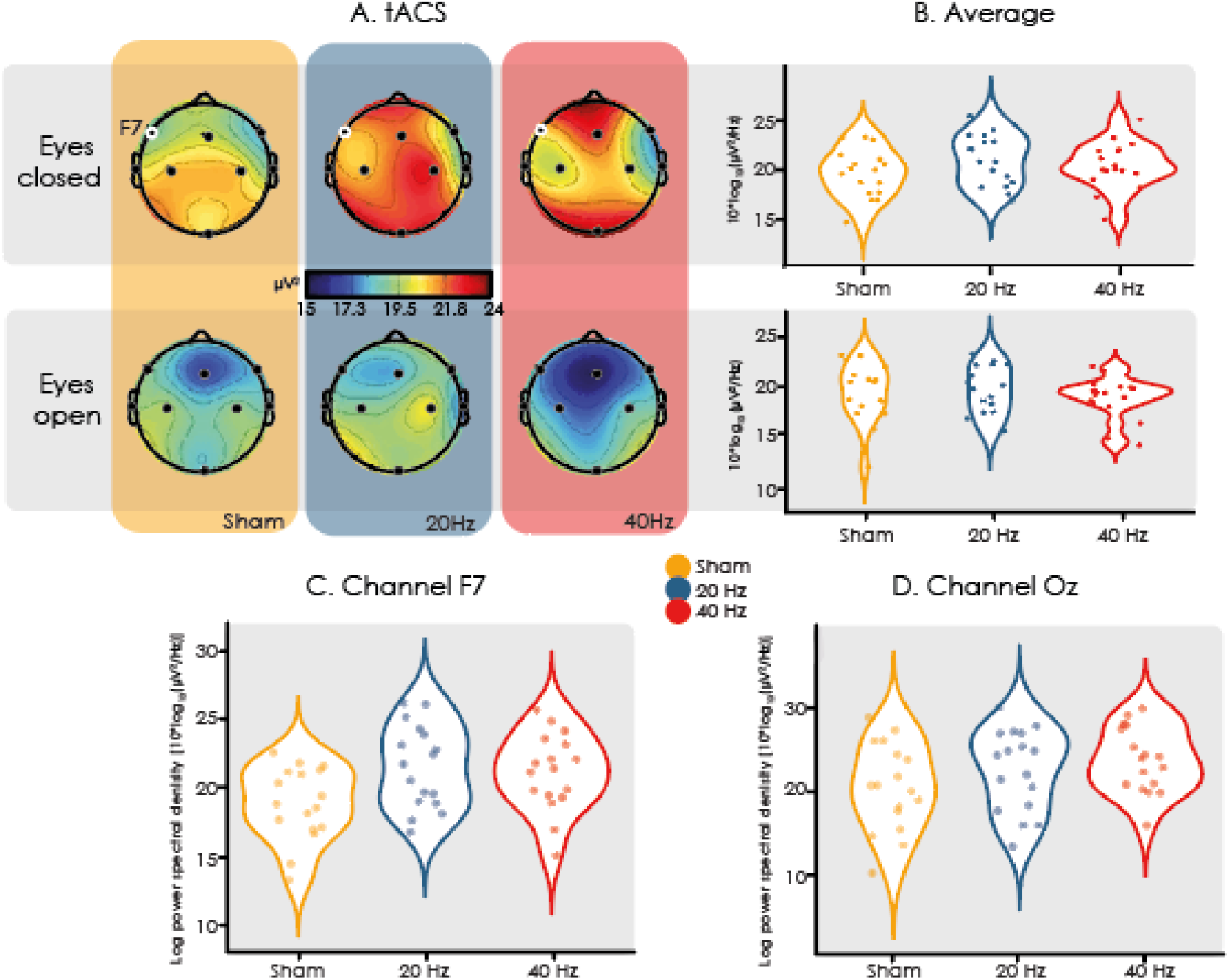
Differences in 20 Hz spectral power between tACS conditions. **a)** Baseline EEG spectral power was subtracted from post tACS EEG. Both 20Hz-tACS and 40Hz-tACS increased 20 Hz power compared to sham-tACS during the eyes closed condition. This increase was only significant for channel F7 (highlighted dot). No significant spectral power differences were found for the eyes open condition. **b)** Differences in log power spectral density between tACS conditions for all averaged electrodes. **c)** Results for frontal channel (F7). Notice that both 20Hz-tACS (t(16) = −3.17, p = 0.048) and 40Hz-tACS (t(16) = −3.17, p = 0.029) show higher 20 Hz power compared to sham-tACS. **d)** Results for occipital central channel (Oz). Increased 20Hz power during the eyes closed condition showed a tendency towards significance (*F*(2,15) =4.99, *p* = 0.0524).

**Figure 6:**
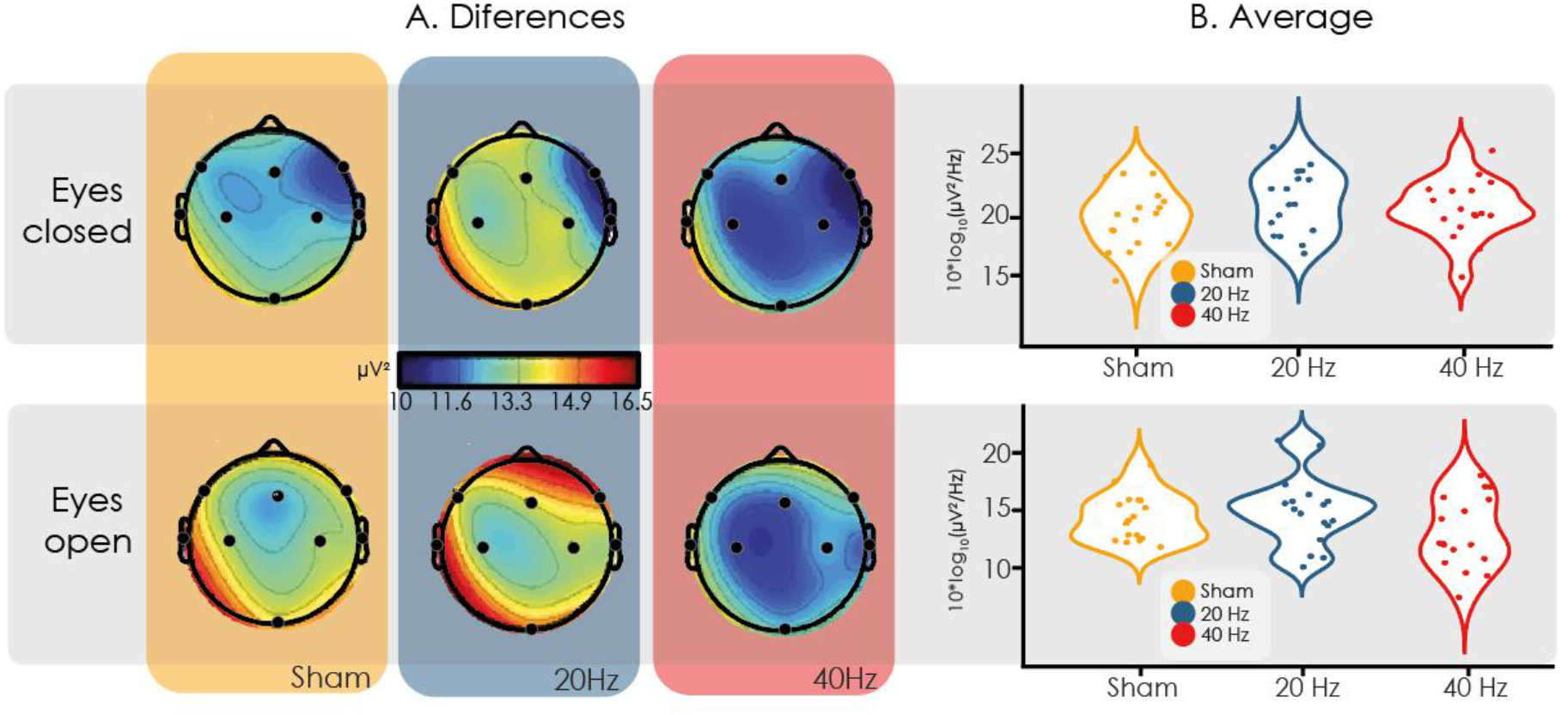
Differences in 40 Hz spectral power between tACS condition. a) Although visual inspection suggests that 40Hz-tACS decreased 40 Hz power both in the eyes open and closed conditions, no statistically significant differences were found. b) Averaged spectral power across all electrodes grouped by tACS stimulation condition.

## Discussion

We implemented a tACS protocol to investigate the possibility of a causal relationship between beta and gamma oscillations and conscious information access. Based on the results of previous experiments and on theoretical considerations, we expected to find that beta-tACS affected the subjective visibility of visual targets, due to the proposed relation between these frequency bands and conscious processes (Lamme and Roelfsema 2000; Engel et al. 2001; Sergent and Dehaene 2004; Melloni et al. 2007; Gaillard et al. 2009; Steinmann et al. 2014; Bastos et al. 2015; Cavinato et al. 2015). Our results confirmed that 20 Hz tACS modulated target visibility, specifically, our data showed that beta-tACS decreased objective visibility and also decreased the blindsight rate (i.e. the amount of ‘correct’ and ‘unseen’ trials) as well as the sightblind rate (i.e. the amount of ‘correct’ and ‘seen’ trials). The blindsight rate is related to subliminal processing, while the sightblind rate is generally related to confidence (Sergent and Dehaene 2004). Since our findings were not specific to either measure, we cannot confirm which brain mechanism was impaired. We also assessed questionnaires to determine potential somatosensory effects of tACS stimulation, and found an increase in discomfort during beta-tACS which could have generally affected the subjects’ perceptual and cognitive capacities. We believe that obtaining reports of “how the experiment itself was experimented” by the participants is paramount when studying subjective experience, and should be generally included in this type of experiments.

The main result of our study is the decrease in objective visibility under beta-tACS stimulation. Before attempting to interpret this result, we first need to discuss the lack of basic mechanistic understanding of how external oscillatory electrical fields interact with neural activity. Multiple reports show that low-frequency tACS applied during NREM in healthy human subjects can enhance associative memory, presumably by entraining these nested rhythms (Marshall et al. 2006; Antonenko et al. 2013; Binder et al. 2014; Prehn-Kristensen et al. 2014; Westerberg et al. 2015; Ladenbauer et al. 2017). Also, there is evidence that tACS applied at the peak oscillatory frequency is capable of entraining these endogenous rhythms in the gamma (Reato et al. 2010; Strüber et al. 2014; Voss et al. 2014), beta (Pogosyan et al. 2009), alpha (Zaehle et al. 2010; Merlet et al. 2013), theta (Marshall et al. 2011), and delta bands (Marshall et al. 2006; Ozen et al. 2010; Fröhlich and McCormick 2010). However, these results have been difficult to replicate, coexisting with several reports of no significant behavioral nor physiological effects (Brignani et al. 2013; Eggert et al. 2013; Horvath et al. 2015a, b; Sahlem et al. 2015). In a recent study, Lafon et al. showed that low-frequency tACS (of 2.5 mA intensity) was unable to modulate spindle and gamma dynamics during NREM sleep (Lafon et al. 2017, 2018). Following a different approach, a recent review claims that stimulation with ~1 mA peak intensity is able to induce only 0.1-0.2 mV changes in membrane potential (Liu et al. 2018). Therefore, the behavioral effects that are induced by tACS are possibly not a result of alterations in neural activity, but of activation restricted to peripheral nerves.

To test whether our findings were actually due to a change in beta power we studied EEG spectral power differences immediately before and after the application of tACS. Our results showed increased beta power after both beta- and gamma-tACS, suggesting that beta power increase could represent a non-specific effect of the stimulation, rather than the result of entrainment of beta oscillations by tACS. A plausible explanation is that tACS could have stimulated subcutaneous nerves, which in turn signaled to the brain and changed EEG power (Liu et al. 2018). It should also be noted that significant beta power differences were only observed in one frontal channel and during the eyes closed condition, which suggests that even if tACS was capable of inducing beta oscillations, this effect might have only been local.

Previous research suggests that the long-range synchrony necessary for conscious access cannot be easily achieved in the high-frequency range (Lachaux et al. 2005; Fries 2005; Buschman and Miller 2007; Buzsáki 2009; Noble and Smith 2015), proposing synchrony in the beta range as the main contributor to this phenomenon. Based on this we would expect facilitated conscious access whenever the neural beta oscillations are promoted and the opposite in those cases where these waves are somehow impaired. Thus, the significant decrease in target visibility under 20 Hz TACS could be explained by the possibility that instead of increasing beta synchrony between distinct brain areas, we interfered with the synchronizing activity. In line with this, Helfrich and colleagues (Helfrich et al. 2014a) showed that while in-phase interhemispheric 40Hz-tACS enhanced parieto-occipital synchronization and the associated perceptual correlates, anti-phase stimulation reduced functional coupling. Also, we cannot rule out the possibility that our tACS protocol only induced small local increases in beta power but was not capable of establishing inter-areal beta synchrony, a possibility that could be investigated using high density EEG combined with source modeling and measures such as phase transfer entropy or granger causality. In future experiments, it might also be interesting to apply closed-loop tACS to modulate brain activity in a phase-dependent manner.

During beta-tACS (compared to sham-tACS), subjects scored higher in their answers to the questionnaire regarding the somatosensory effects of the stimulation, such as tickling, burning or visual light flickering. This adds support to the possibility that behavioral differences were ultimately the result of non-neural effects of tACS. For example, the somatosensory effects could have distracted the subjects, resulting in worse performance. However, as previously noted, no correlation was found between the questionnaire scores and performances. Importantly, while this kind of questionnaire is seldomly used in studies using tACS or other forms of stimulation, we have shown that it can provide useful information for the interpretation of the experimental results. Our analysis of the questionnaires demonstrated significantly higher somatosensory scores for the 20Hz-tACS condition, consistent with a previous study that used a similar questionnaire and applied the same stimulation intensity, establishing that tACS-induced cutaneous sensations are frequency-dependent and are most perceptible at 20 Hz alternating stimulation (Turi et al. 2013). Since multiple studies concluded that subjects are not capable of distinguishing active vs. sham tACS, we are not concerned with the possibility of unblinding (Wach et al. 2013a, b). Instead, our concern lies in the potential effects of sensory intrusion during the performance of the task. To assess this possibility, we correlated the questionnaire answers with the target visibility modulation and did not find a significant association between both variables.

We emphasize that our work showed that beta-tACS but not gamma-tACS affected visual perception, i.e. that our findings are frequency-specific. Of note, the majority of EEG studies on the neural correlates of conscious information access focus on gamma instead of on beta oscillations. In contrast with this trend, our work established the modulation of target visibility by beta- and not gamma-tACS. The fact that numerous studies showed increased gamma power and synchrony during the resolving of ambiguous or bistable stimuli (Laczó et al. 2012; Helfrich et al. 2014a; Cabral-Calderin et al. 2015) might be because gamma oscillations are important for feature binding rather than for conscious information access. Nunez & Srinivasan (2010) already suggested that gamma band oscillations are overly emphasized in studies of conscious perception. Lower frequency oscillations, such as beta, capable of synchronizing large neural populations over great distances, in contrast to gamma oscillations (Kopell et al. 2000; von Stein and Sarnthein 2000; Buzsáki and Draguhn 2004). Importantly, studies of the visual cortex showed that gamma oscillations travel in a feedforward direction, while lower frequencies, such as beta and alpha, travel in a feedback direction (van Kerkoerle et al. 2014; Bastos et al. 2015), and thus they might play a fundamental role in conscious perception, insofar it necessitates top-bottom feedback for the amplification of the incoming sensory information.

Going beyond the dichotomy between beta and gamma oscillations, a complete assessment of these and other findings will inevitably highlight the complexity of neural oscillations as a candidate mechanism for inter-areal communication in the brain, leading to the conclusion that the details of visual processing and conscious information access go beyond their characterization in terms of band-specific feedforward or feedback processes. There is a marked tendency to associate well-defined cognitive functions to specific oscillatory frequencies; however, it is unlikely that oscillations can be mapped into completely separate functions. Moreover, it has been shown that different frequencies present mutual interactions, such as when fast frequencies are nested on slower ones (Roopun et al. 2008; Belluscio et al. 2012). In this case, the identification of feedforward and feedback activity with gamma and beta oscillations, respectively, might constitute an overly simplistic model. Even if this is not the case, strong mutual interdependences between cortical oscillations could impede the selective empirical manipulation of narrow frequency bands by external means, as we attempted in the present study.

An intriguing finding is that we did not observe the nonlinear increase in objective and subjective visibility as a function of SOA, as shown by Del Cul and colleagues (Del Cul et al. 2007). An important difference between the task we adopted for our this study and that used by Del Cul et al. is that we included a personalized target contrast obtained in the staircase session. The objective of including a personalized contrast was to control for inter-subject variability, setting a common visibility threshold at a SOA of 50 ms. However, the resulting visibility threshold was considerably lower, therefore, it might be possible that on average the contrast was too high and consequently that the task was too easy for the participants. In this case, the sudden increase in visibility would not be found because the target numerals were already visible at the lower SOAs. Another possibility is that conscious perception can occur more gradually than anticipated. The GW theory posits that the neural basis of conscious access consists of an all-or-none self-amplifying process, which results in a global and persisting pattern of activity (Sergent and Dehaene 2004). In contrast, other researchers proposed that conscious perception is a continuum, associated with a gradual change in the intensity of brain activation, until reaching a certain level at which sensory information becomes consciously accessible (Nieuwenhuis and de Kleijn 2011). Theoretical considerations aside, the way the subjects understand the instructions to report and judge what constitutes conscious perception for them is likely to influence the nonlinearity of the subjective visibility curve. As in the case of the questionnaire to assess the somatosensory effects of tACS, a careful debriefing of the participants concerning the details of their experiences during the backward masking paradigm could be useful to resolve this apparent discrepancy.

In summary, this study showed that beta-tACS is capable of modulating target visibility in a metacontrast backward-masking task. While interesting and novel, it is not yet certain that this modulation is mediated by an effect of tACS on neural activity. To add support to this possibility, future studies should investigate inter-areal connectivity changes as a response to beta-tACS. At the time of this study, the tACS literature presents some important contradictions: recent research suggests that higher intensities than those generally applied (2.5 mA) are necessary for the entrainment of neural oscillations, yet entrainment is frequently cited as the principal mechanism of action of tACS, regardless of this potential issue. However, it could be possible to induce “resonant” effects of tACS on intrinsic brain activity by means of in-phase stimulation delivered via closed-loop setups (Ketz et al. 2018), a possibility to be explored in future experiments. Finally, even under optimal conditions, the selective manipulation of a frequency band by non-invasive electrical stimulation could be difficult or even impossible, highlighting the need to back up our conclusions with further work in animal models, in combination with computational studies capable of exploring the mechanistic causes and implications of our findings

## Supplementary results

**Supplementary figure 1:**
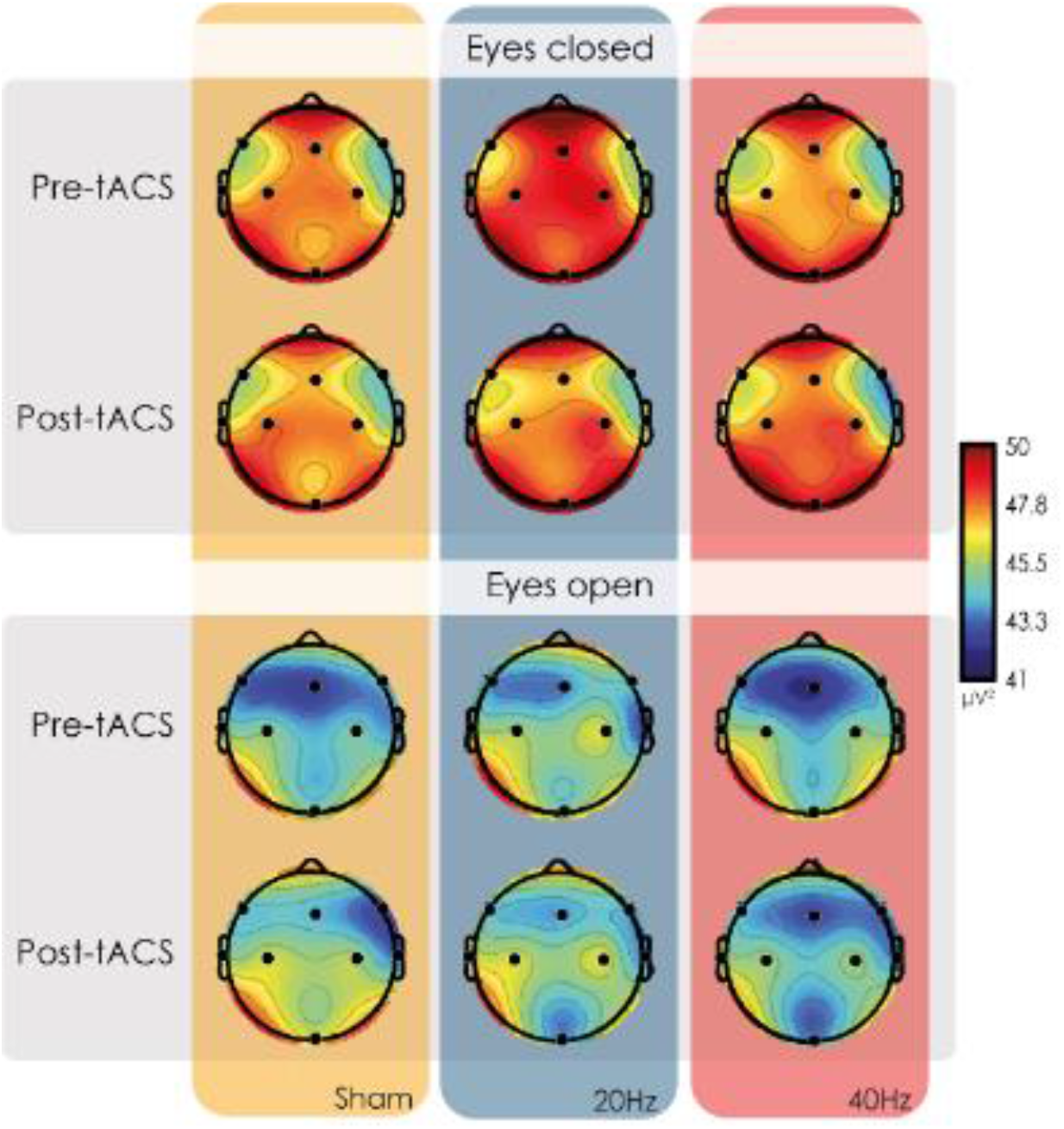
Same EEG topographies as in figure 5 (20Hz power differences) but without subtracting baseline EEG (pre-tACS) from post-tACS EEG.

**Supplementary figure 2:**
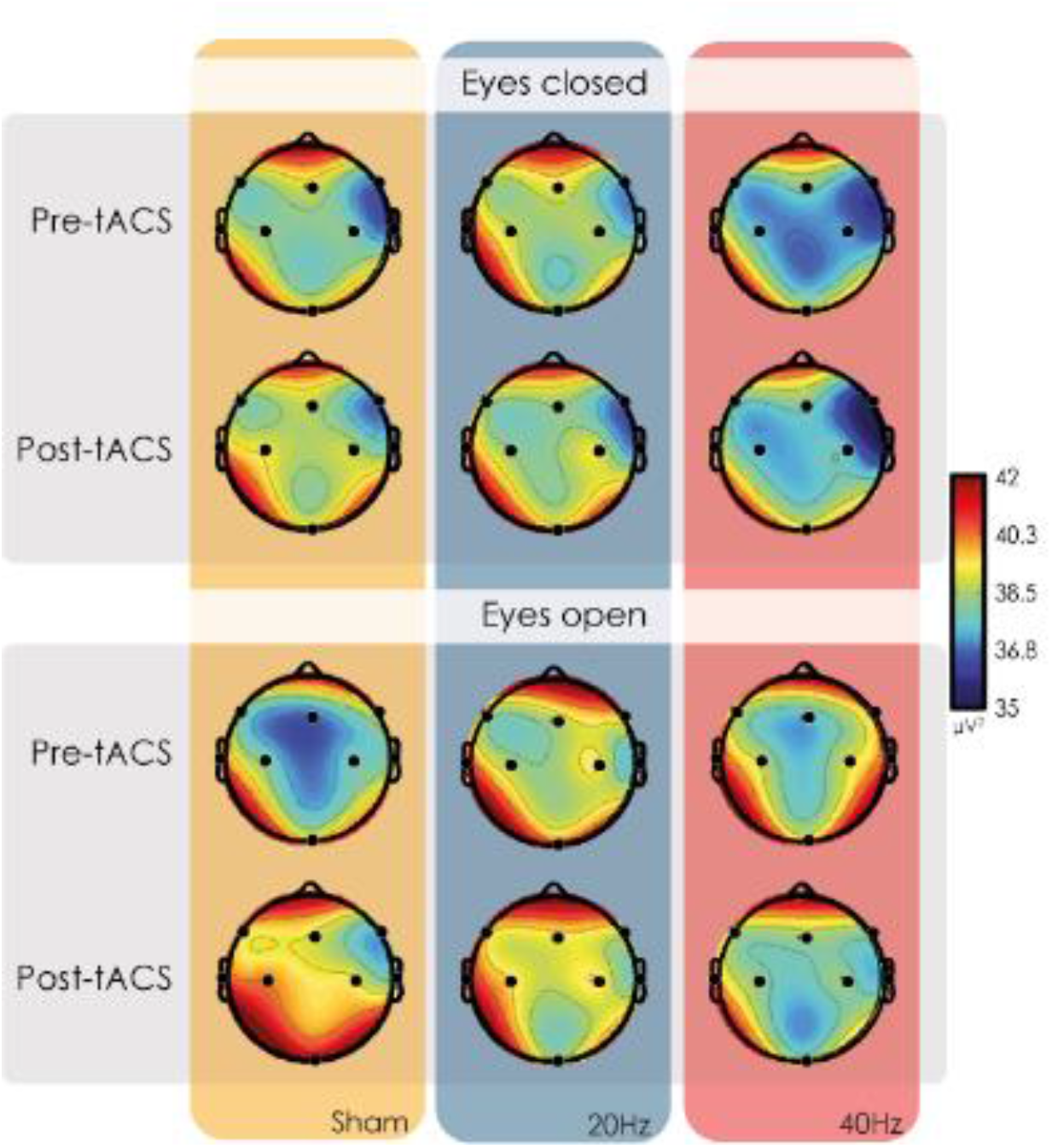
Same EEG topographies as in figure 6 (40Hz power differences) but without subtracting baseline EEG (pre-tACS) from post-tACS EEG.

## References

Antal A, Alekseichuk I, Bikson M, et al (2017) Low intensity transcranial electric stimulation: Safety, ethical, legal regulatory and application guidelines. Clin Neurophysiol 128:1774–1809. https://doi.org/10.1016/J.CLINPH.2017.06.001

Antonenko D, Diekelmann S, Olsen C, et al (2013) Napping to renew learning capacity: Enhanced encoding after stimulation of sleep slow oscillations. Eur J Neurosci 37:1142–1151. https://doi.org/10.1111/ejn.12118

Baars BJ (2002) The conscious access hypothesis: origins and recent evidence. Trends Cogn Sci 6:47–52. https://doi.org/10.1016/S1364-6613(00)01819-2

Baars BJ (2005) Global workspace theory of consciousness: toward a cognitive neuroscience of human experience. Prog Brain Res 150:45–53. https://doi.org/10.1016/S0079-6123(05)50004-9

Baars BJ, Franklin S, Ramsoy TZ (2013) Global Workspace Dynamics: Cortical “Binding and Propagation” Enables Conscious Contents. Front Psychol 4:200. https://doi.org/10.3389/fpsyg.2013.00200

Bastos AM, Vezoli J, Bosman CA, et al (2015) Visual Areas Exert Feedforward and Feedback Influences through Distinct Frequency Channels. Neuron 85:390–401. https://doi.org/10.1016/J.NEURON.2014.12.018

Belluscio MA, Mizuseki K, Schmidt R, et al (2012) Cross-Frequency Phase-Phase Coupling between Theta and Gamma Oscillations in the Hippocampus. J Neurosci 32(2):423–435. https://doi.org/10.1523/JNEUROSCI.4122-11.2012

Binder S, Berg K, Gasca F, et al (2014) Transcranial slow oscillation stimulation during sleep enhances memory consolidation in rats. Brain Stimul 7:508–515. https://doi.org/10.1016/j.brs.2014.03.001

Bliksted V, Samuelsen E, Sandberg K, et al (2017) Discriminating between first- and second-order cognition in first-episode paranoid schizophrenia. Cogn Neuropsychiatry 22:95–107. https://doi.org/10.1080/13546805.2016.1268954

Brignani D, Ruzzoli M, Mauri P, Miniussi C (2013) Is Transcranial Alternating Current Stimulation Effective in Modulating Brain Oscillations? PLoS One 8:. https://doi.org/10.1371/journal.pone.0056589

Buschman TJ, Miller EK (2007) Top-down versus bottom-up control of attention in the prefrontal and posterior parietal cortices. Science (80-) 315:1860–1864. https://doi.org/10.1126/science.1138071

Buzsáki G (2009) Rhythms of the Brain

Buzsáki G, Draguhn A (2004) Neuronal oscillations in cortical networks. Science 304:1926–9. https://doi.org/10.1126/science.1099745

Cabral-Calderin Y, Schmidt-Samoa C, Wilke M (2015) Rhythmic Gamma Stimulation Affects Bistable Perception. J Cogn Neurosci 27:1298–1307. https://doi.org/10.1162/jocn_a_00781

Cavinato M, Genna C, Manganotti P, et al (2015) Coherence and Consciousness: Study of Fronto-Parietal Gamma Synchrony in Patients with Disorders of Consciousness. Brain Topogr 28:570–579. https://doi.org/10.1007/s10548-014-0383-5

Chang M, Iizuka H, Kashioka H, et al (2017) Unconscious improvement in foreign language learning using mismatch negativity neurofeedback: A preliminary study. PLoS One 12:e0178694. https://doi.org/10.1371/journal.pone.0178694

Crick F, Koch C] (1990) Towards a neurobiological theory of consciousness

Dehaene S, Changeux J-P, Naccache L (2011) The Global Neuronal Workspace Model of Conscious Access: From Neuronal Architectures to Clinical Applications. Springer, Berlin, Heidelberg, pp 55–84

Dehaene S, Changeux JP, Naccache L, et al (2006) Conscious, preconscious, and subliminal processing: a testable taxonomy. Trends Cogn Sci 10:204–211. https://doi.org/10.1016/j.tics.2006.03.007

Dehaene S, Naccache L (2001) Towards a cognitive neuroscience of consciousness: basic evidence and a workspace framework. Cognition 79:1–37. https://doi.org/10.1016/S0010-0277(00)00123-2

Dehaene S, Naccache L, Cohen L, et al (2001) Cerebral mechanisms of word masking and unconscious repetition priming. Nat Neurosci 4:752–758. https://doi.org/10.1038/89551

Dehaene S, Naccache L, Le Clec’H G, et al (1998) Imaging unconscious semantic priming. Nature 395:597–600. https://doi.org/10.1038/26967

Del Cul A, Baillet S, Dehaene S (2007) Brain Dynamics Underlying the Nonlinear Threshold for Access to Consciousness. PLoS Biol 5:e260. https://doi.org/10.1371/journal.pbio.0050260

Eggert T, Dorn H, Sauter C, et al (2013) No effects of slow oscillatory transcranial direct current stimulation (tDCS) on sleep-dependent memory consolidation in healthy elderly subjects. Brain Stimul 6:938–945. https://doi.org/10.1016/j.brs.2013.05.006

Engel AK, Fries P, Singer W (2001) Dynamic predictions: oscillations and synchrony in top-down processing. Nat Rev Neurosci 2:704–716

Engel AK, Singer W (2001) Temporal binding and the neural correlates of sensory awareness. Trends Cogn Sci 5:16–25. https://doi.org/10.1016/S1364-6613(00)01568-0

Fries P (2005) A mechanism for cognitive dynamics: neuronal communication through neuronal coherence. Trends Cogn Sci 9:474–80. https://doi.org/10.1016/j.tics.2005.08.011

Fröhlich F, McCormick DA (2010) Endogenous Electric Fields May Guide Neocortical Network Activity. Neuron 67:129–143. https://doi.org/10.1016/j.neuron.2010.06.005

Gaillard R, Dehaene S, Adam C, et al (2009) Converging Intracranial Markers of Conscious Access. PLoS Biol 7:e1000061. https://doi.org/10.1371/journal.pbio.1000061

Helfrich RF, Knepper H, Nolte G, et al (2014a) Selective Modulation of Interhemispheric Functional Connectivity by HD-tACS Shapes Perception. PLoS Biol 12:e1002031. https://doi.org/10.1371/journal.pbio.1002031

Helfrich RF, Schneider TR, Rach S, et al (2014b) Entrainment of Brain Oscillations by Transcranial Alternating Current Stimulation. Curr Biol 24:333–339. https://doi.org/10.1016/J.CUB.2013.12.041

Horvath JC, Forte JD, Carter O (2015a) Quantitative review finds no evidence of cognitive effects in healthy populations from single-session transcranial direct current stimulation (tDCS). Brain Stimul 8:535–550. https://doi.org/10.1016/j.brs.2015.01.400

Horvath JC, Forte JD, Carter O (2015b) Evidence that transcranial direct current stimulation (tDCS) generates little-to-no reliable neurophysiologic effect beyond MEP amplitude modulation in healthy human subjects: A systematic review. Neuropsychologia 66:213–236. https://doi.org/10.1016/j.neuropsychologia.2014.11.021

Ketz N, Jones AP, Bryant NB, et al (2018) Closed-Loop Slow-Wave tACS Improves Sleep-Dependent Long-Term Memory Generalization by Modulating Endogenous Oscillations. J Neurosci 38:7314–7326. https://doi.org/10.1523/JNEUROSCI.0273-18.2018

Kopell N, Ermentrout GB, Whittington MA, Traub RD (2000) Gamma rhythms and beta rhythms have different synchronization properties. Proc Natl Acad Sci U S A 97:1867–72. https://doi.org/10.1073/pnas.97.4.1867

Lachaux JP, George N, Tallon-Baudry C, et al (2005) The many faces of the gamma band response to complex visual stimuli. Neuroimage 25:491–501. https://doi.org/10.1016/j.neuroimage.2004.11.052

Laczó B, Antal A, Niebergall R, et al (2012) Transcranial alternating stimulation in a high gamma frequency range applied over V1 improves contrast perception but does not modulate spatial attention. Brain Stimul 5:484–491. https://doi.org/10.1016/J.BRS.2011.08.008

Ladenbauer J, Ladenbauer J, Külzowä N, et al (2017) Promoting sleep oscillations and their functional coupling by transcranial stimulation enhances memory consolidation in mild cognitive impairment. J Neurosci 37:7111–7124. https://doi.org/10.1523/JNEUROSCI.0260-17.2017

Lafon B, Henin S, Huang Y, et al (2017) Low frequency transcranial electrical stimulation does not entrain sleep rhythms measured by human intracranial recordings. Nat Commun 8:1–14. https://doi.org/10.1038/s41467-017-01045-x

Lafon B, Henin S, Huang Y, et al (2018) Erratum: Author Correction: Low frequency transcranial electrical stimulation does not entrain sleep rhythms measured by human intracranial recordings (Nature communications (2017) 8 1 (1199)). Nat Commun 9:949. https://doi.org/10.1038/s41467-018-03392-9

Lamme VAF, Roelfsema PR (2000) The distinct modes of vision offered by feedforward and recurrent processing. Trends Neurosci 23:571–579. https://doi.org/10.1016/S0166-2236(00)01657-X

Liu A, Vöröslakos M, Kronberg G, et al (2018) Immediate neurophysiological effects of transcranial electrical stimulation. Nat Commun 9:5092. https://doi.org/10.1038/s41467-018-07233-7

Łukowska M, Sznajder M, Wierzchoń M (2018) Error-related cardiac response as information for visibility judgements. Sci Reports 2018 81 8:1131. https://doi.org/10.1038/s41598-018-19144-0

Marshall L, Helgadóttir H, Mölle M, Born J (2006) Boosting slow oscillations during sleep potentiates memory. Nature 444:610–613. https://doi.org/10.1038/nature05278

Marshall L, Kirov R, Brade J, et al (2011) Transcranial electrical currents to probe EEG brain rhythms and memory consolidation during sleep in humans. PLoS One 6:. https://doi.org/10.1371/journal.pone.0016905

Melloni L, Molina C, Pena M, et al (2007) Synchronization of neural activity across cortical areas correlates with conscious perception. J Neurosci 27:2858–2865

Merlet I, Birot G, Salvador R, et al (2013) From Oscillatory Transcranial Current Stimulation to Scalp EEG Changes: A Biophysical and Physiological Modeling Study. PLoS One 8:1–12. https://doi.org/10.1371/journal.pone.0057330

Naccache L, Dehaene S (2001) The Priming Method: Imaging Unconscious Repetition Priming Reveals an Abstract Representation of Number in the Parietal Lobes. Cereb Cortex 11:966–974. https://doi.org/10.1093/cercor/11.10.966

Naro A, Bramanti P, Leo A, et al (2016) Transcranial Alternating Current Stimulation in Patients with Chronic Disorder of Consciousness: A Possible Way to Cut the Diagnostic Gordian Knot? Brain Topogr 29:623–644. https://doi.org/10.1007/s10548-016-0489-z

Nieuwenhuis S, de Kleijn R (2011) Consciousness of targets during the attentional blink: a gradual or all-or-none dimension? Attention, Perception, Psychophys 73:364–373. https://doi.org/10.3758/s13414-010-0026-1

Noble H, Smith J (2015) untitled _ Enhanced Reader.pdf. Clin. Infect. Dis.

Nunez PL, Srinivasan R (2010) Scale and frequency chauvinism in brain dynamics: too much emphasis on gamma band oscillations. Brain Struct Funct 215:67–71. https://doi.org/10.1007/s00429-010-0277-6

Ozen S, Sirota A, Belluscio MA, et al (2010) Transcranial electric stimulation entrains cortical neuronal populations in rats. J Neurosci 30:11476–11485. https://doi.org/10.1523/JNEUROSCI.5252-09.2010

Peirce JW, Gray JR, Simpson S, et al (2019) PsychoPy2: experiments in behavior made easy.

Pogosyan A, Gaynor LD, Eusebio A, Brown P (2009) Boosting Cortical Activity at Beta-Band Frequencies Slows Movement in Humans. Curr. Biol. 19:1637–1641

Prehn-Kristensen A, Munz M, Göder R, et al (2014) Transcranial oscillatory direct current stimulation during sleep improves declarative memory consolidation in children with attention-deficit/hyperactivity disorder to a level comparable to healthy controls. Brain Stimul 7:793–799. https://doi.org/10.1016/j.brs.2014.07.036

R Core Team (2014) R: A language and environment for statistical computing.

Reato D, Rahman A, Bikson M, Parra LC (2010) Low-intensity electrical stimulation affects network dynamics by modulating population rate and spike timing. J Neurosci 30:15067–15079. https://doi.org/10.1523/JNEUROSCI.2059-10.2010

Roopun AK, Kramer MA, Carracedo LM, et al (2008) concatenation underlies interactions between gamma and beta rhythms in neocortex. Front Cell Neurosci 2:1. https://doi.org/10.3389/neuro.03.001.2008

Sahlem GL, Badran BW, Halford JJ, et al (2015) Oscillating Square Wave Transcranial Direct Current Stimulation (tDCS) Delivered During Slow Wave Sleep Does Not Improve Declarative Memory More Than Sham: A Randomized Sham Controlled Crossover Study. Brain Stimul 8:528–534. https://doi.org/10.1016/j.brs.2015.01.414

Sergent C, Dehaene S (2004) Neural processes underlying conscious perception: Experimental findings and a global neuronal workspace framework. J Physiol 98:374–384. https://doi.org/10.1016/J.JPHYSPARIS.2005.09.006

Seth AK, Dienes Z, Cleeremans A, et al (2008) Measuring consciousness: relating behavioural and neurophysiological approaches. Trends Cogn Sci 12:314–321. https://doi.org/10.1016/j.tics.2008.04.008

Steinmann S, Leicht G, Ertl M, et al (2014) Conscious auditory perception related to long-range synchrony of gamma oscillations. Neuroimage 100:435–443. https://doi.org/10.1016/J.NEUROIMAGE.2014.06.012

Strüber D, Rach S, Trautmann-Lengsfeld SA, et al (2014) Antiphasic 40 Hz oscillatory current stimulation affects bistable motion perception. Brain Topogr 27:158–171. https://doi.org/10.1007/s10548-013-0294-x

Tagliazucchi E (2020) Consciousness and Its Disorders. In: Reference Module in Neuroscience and Biobehavioral Psychology. Elsevier, pp 1–12

Turi Z, Ambrus GG, Janacsek K, et al (2013) Both the cutaneous sensation and phosphene perception are modulated in a frequency-specific manner during transcranial alternating current stimulation. Restor Neurol Neurosci 31:275–285. https://doi.org/10.3233/RNN-120297

van Gaal S, Naccache L, Meuwese JDI, et al (2014) Can the meaning of multiple words be integrated unconsciously? Philos Trans R Soc B Biol Sci 369:20130212–20130212. https://doi.org/10.1098/rstb.2013.0212

van Kerkoerle T, Self MW, Dagnino B, et al (2014) Alpha and gamma oscillations characterize feedback and feedforward processing in monkey visual cortex. Proc Natl Acad Sci U S A 111:14332–41. https://doi.org/10.1073/pnas.1402773111

von Stein A, Sarnthein J (2000) Different frequencies for different scales of cortical integration: from local gamma to long range alpha/theta synchronization. Int J Psychophysiol 38:301–313. https://doi.org/10.1016/S0167-8760(00)00172-0

Voss U, Holzmann R, Hobson A, et al (2014) Induction of self awareness in dreams through frontal low current stimulation of gamma activity. Nat Neurosci 17:810–812. https://doi.org/10.1038/nn.3719

Vossen A, Gross J, Thut G (2015) Alpha Power Increase After Transcranial Alternating Current Stimulation at Alpha Frequency (α-tACS) Reflects Plastic Changes Rather Than Entrainment. Brain Stimul 8:499–508. https://doi.org/10.1016/J.BRS.2014.12.004

Wach C, Krause V, Moliadze V, et al (2013a) Effects of 10 Hz and 20 Hz transcranial alternating current stimulation (tACS) on motor functions and motor cortical excitability. Behav Brain Res 241:1–6. https://doi.org/10.1016/J.BBR.2012.11.038

Wach C, Krause V, Moliadze V, et al (2013b) The effect of 10 Hz transcranial alternating current stimulation (tACS) on corticomuscular coherence. Front Hum Neurosci 7:511. https://doi.org/10.3389/fnhum.2013.00511

Westerberg CE, Florczak SM, Weintraub S, et al (2015) Memory improvement via slow-oscillatory stimulation during sleep in older adults. Elsevier Ltd

Zaehle T, Rach S, Herrmann CS (2010) Transcranial Alternating Current Stimulation Enhances Individual Alpha Activity in Human EEG. PLoS One 5:1–7. https://doi.org/10.1371/journal.pone.0013766

(2018) The MathWorks

